# Maternal manipulation of offspring size can trigger the evolution of eusociality in promiscuous species

**DOI:** 10.1101/2024.01.23.576864

**Authors:** Ella Rees-Baylis, Ido Pen, Jan J. Kreider

**Affiliations:** Theoretical Research in Evolutionary Life Sciences, Groningen Institute for Evolutionary Life Sciences, University of Groningen, Nijenborgh 7, 9747 AG Groningen, The Netherlands

**Author notes:** Corresponding author JJK. Equally contributing authors (Ido Pen, Jan J. Kreider).

## Abstract

Eusocial organisms typically live in colonies with one reproductive queen supported by thousands of sterile workers. It is widely believed that monogamous mating is a precondition for the evolution of eusociality. Here, we present a theoretical model that simulates a realistic scenario for the evolution of eusociality. In the model, mothers can evolve control over resource allocation to offspring, affecting offspring body size. The offspring can evolve body-size-dependent dispersal, by which they disperse to breed or stay at the nest as helpers. We demonstrate that eusociality evolves, even if mothers are not strictly monogamous, if mothers can constrain their offspring’s reproduction by manipulation. We also observe the evolution of social polymorphism with small individuals that help and larger individuals that disperse to breed. Our model unifies the traditional kin selection and maternal manipulation explanations for the evolution of eusociality and demonstrates that – contrary to current consensus belief – eusociality can evolve despite highly promiscuous mating.

## Introduction

Reproductive altruism, where individuals forfeit their own reproduction by committing to non-reproductive helper roles, has evolved independently multiple times across animal societies^1,2^. For instance, in naked mole rats, a sole female breeds whilst the remaining females perform nest building and foraging^3,4^. In termites, ants, some bees, and some wasps, the queen is supported by numerous smaller workers that may not reproduce, instead partaking in foraging, nest defence, and care for young^5–7^. Explaining the evolution of such eusocial breeding is a core issue of evolutionary biology, since sterile helpers do not reproduce – a behaviour that should be selected against^8–10^.

Natural selection favours individuals to be reproductively altruistic if the inclusive fitness costs of forgoing their own reproduction are outweighed by the inclusive fitness benefits of reproductive altruism^11,12^. Such benefits can be gained if the reproductively altruistic individual (the worker) can enhance the success of genes shared with the beneficiary of the altruistic behaviour (the queen) through behaving altruistically. Thus, high genetic relatedness between the beneficiary and the helping individual should make the evolution of reproductive altruism and eusociality more likely^9,10^. It has therefore been proposed that lifetime monogamy of the breeding female is an essential prerequisite for the evolution of eusociality, as it enables newly emerged offspring to help raise their full siblings, to whom they are highly related^13–16^.

Another essential requirement for the evolution of eusociality is the presence of overlapping generations to allow offspring to care for their siblings from the subsequent generation^17,18^. The simplest form of such overlapping generations is an annual life cycle where a breeding female produces two broods per year, enabling offspring from the first brood to help raise their siblings from the second brood. This life cycle – a partially bivoltine life cycle – is found in many non-eusocial species of bees and wasps that are closely related to eusocial species, and it is hypothesised to be ancestral to the evolution of eusociality^19–22^.

Partial bivoltinism can facilitate the evolution of eusociality because breeding females can temporally split offspring sex ratios across broods. This favours the evolution of reproductive altruism in haplodiploid organisms (e.g. ants, bees and wasps) if females from the first brood are primarily raising their sisters to whom they are more closely related than to their brothers^23,24^. The two distinct broods of a partially bivoltine life cycle also enable pre-existing morphological and behavioural differences between broods to be co-opted for the evolution of worker- and queen-phenotypes^20,25,26^. Such pre-existing differences could be based on differential maternal resource allocation strategies between broods. For instance, in some bees, mothers differentially allocate food to their daughters, causing some smaller daughters to remain at the natal nest being coerced into helping^27–30^.

Kin selection and maternal manipulation have often been regarded as alternative explanations for the evolution of eusociality^31–33^, but they are not mutually exclusive^34–36^. Here, we present an evolutionary individual-based model to unify kin selection and maternal manipulation explanations for the evolution of eusociality whilst also explicitly modelling phenotypic evolution of queens and workers. We model a partially bivoltine population of haplodiploid organisms (Fig. 1). Female offspring from the first brood evolve a body-size-dependent dispersal strategy to remain at the natal nest as a helper or to disperse and breed independently. The breeding female, in turn, evolves a resource allocation strategy through which she has control over offspring body size. The simulations start with solitary populations; thus, initially offspring disperse and mothers produce offspring that all have identical body sizes. However, if mothers evolve to produce smaller offspring and if smaller offspring have low breeding success, e.g., due to insufficient energy reserves to develop their ovaries, then the co-evolution of maternal resource allocation strategy and body-size-dependent offspring dispersal could lead to the evolution of eusociality, the production of small workers, and thus to queen-worker dimorphism.

**Fig. 1.**
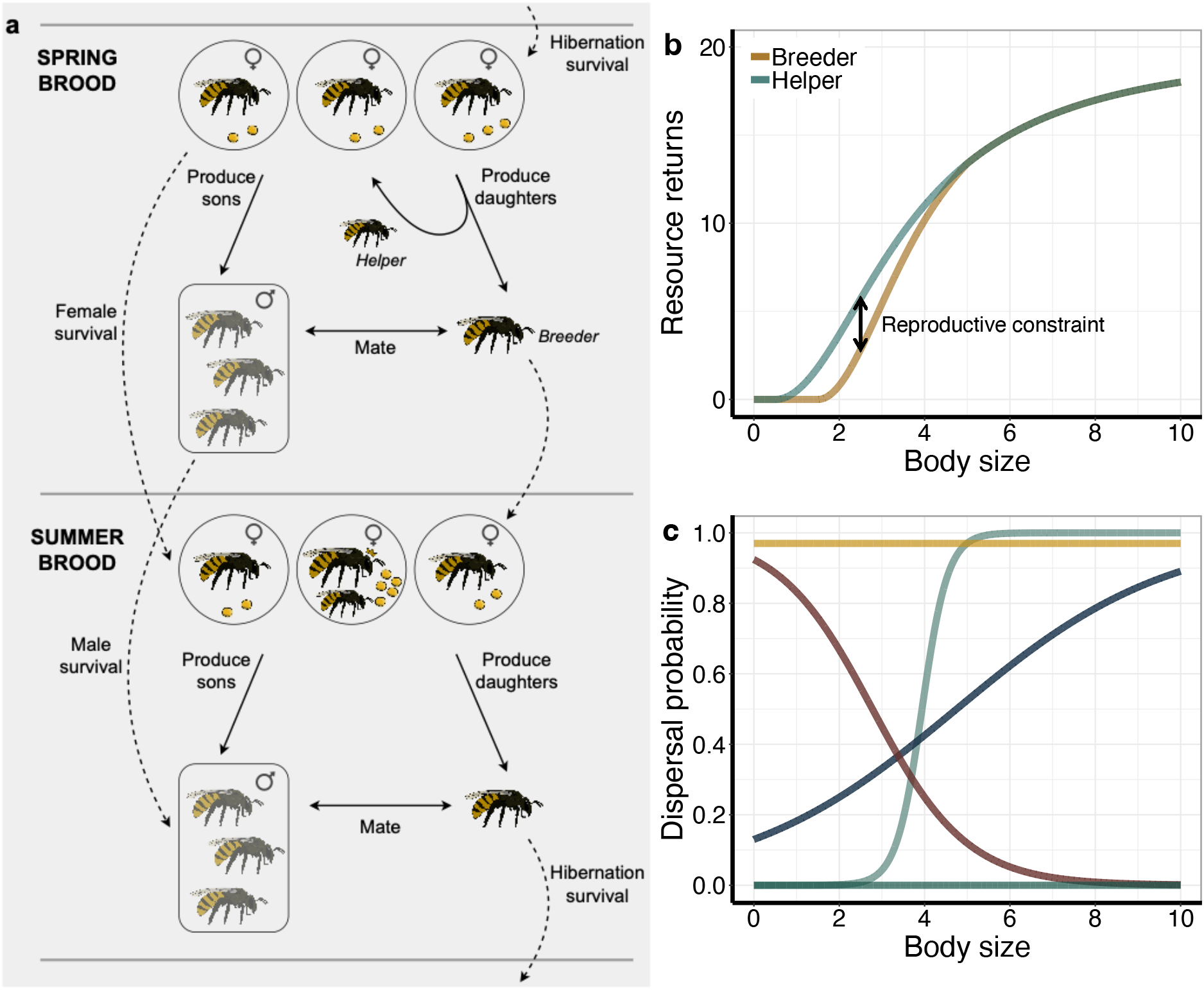
Life cycle, resource return functions, and possible dispersal reaction norms in the model. **(a)** A partially bivoltine life cycle consists of two reproductive periods per year – the spring (top) and summer brood (bottom). In spring, solitary foundresses acquire resources (yellow circles) which they use to produce offspring. Female spring-brood offspring evolve a body-size-dependent dispersal probability, which determines whether they mate, disperse and breed or remain at the natal nest as a helper. During the summer brood, nests can either be solitary, including a surviving foundress or a spring-brood female that dispersed, or they are eusocial if the foundress has at least one helper. Females again acquire resources, with eusocial nests gaining resources by both the breeder and helper(s), and produce offspring. The female summer-brood offspring mates with males from the summer brood or surviving spring-brood males. All males and breeding and helping females die at the end of summer. The female summer-brood offspring hibernate to become solitary foundresses during the following spring. **(b)** The amount of resources gained by a female is dependent on her body size. Small individuals do not acquire any resources. At large body sizes, resource returns diminish. In some model scenarios, we assume a body-size-specific reproductive constraint, rendering small individuals more successful as helpers than they were as breeders. **(c)** Examples of dispersal reaction norms, showing possible evolutionary outcomes for the relationship between dispersal probability and body size of an individual. All females are initiated with the yellow reaction norm; thus, the populations are initially preliminarily solitarily breeding (unless stated otherwise).

## Results

### Partial bivoltinism favours the evolution of eusociality, even under low levels of polyandry

First, we simulated the model by assuming that both breeders and helpers have the same resource return functions (the ‘helper’ function in Fig. 1b now applies to both breeders and helpers). We vary the mating frequency from 1.0 to 2.0 to investigate the effect of polyandrous mating on the evolution of eusociality. As an example, a mating frequency of 1.3 could result from 30% of the females in the population mating with two males and 70% with one male. Assuming identical resource return functions for breeders and helpers, eusociality evolves below a mating frequency of 1.3. At 1.3 mating frequency, only some nests become eusocial. At mating frequencies higher than 1.3, the populations remain solitary. Consequently, strict lifetime monandry is not a necessary requirement for the evolution of eusociality from a partially bivoltine life cycle. This is because male generational overlap in a partially bivoltine life cycle decreases the reproductive value of summer-brood males, thus enabling spring-brood females to capitalise on the relatedness asymmetries from haplodiploidy between their sisters and brothers from the summer brood^24^. Maternal control of body size has no impact on the evolution of eusociality, if the resource returns for breeders and helpers are identical (Fig. 2a). Although resource returns depend on body size, fitness benefits or costs of helping are independent of body size, providing no incentive for spring-females to evolve a body-size-specific dispersal strategy. This demonstrates that maternal control of offspring body size alone is not sufficient for the evolution of eusociality through maternal manipulation.

**Fig. 2.**
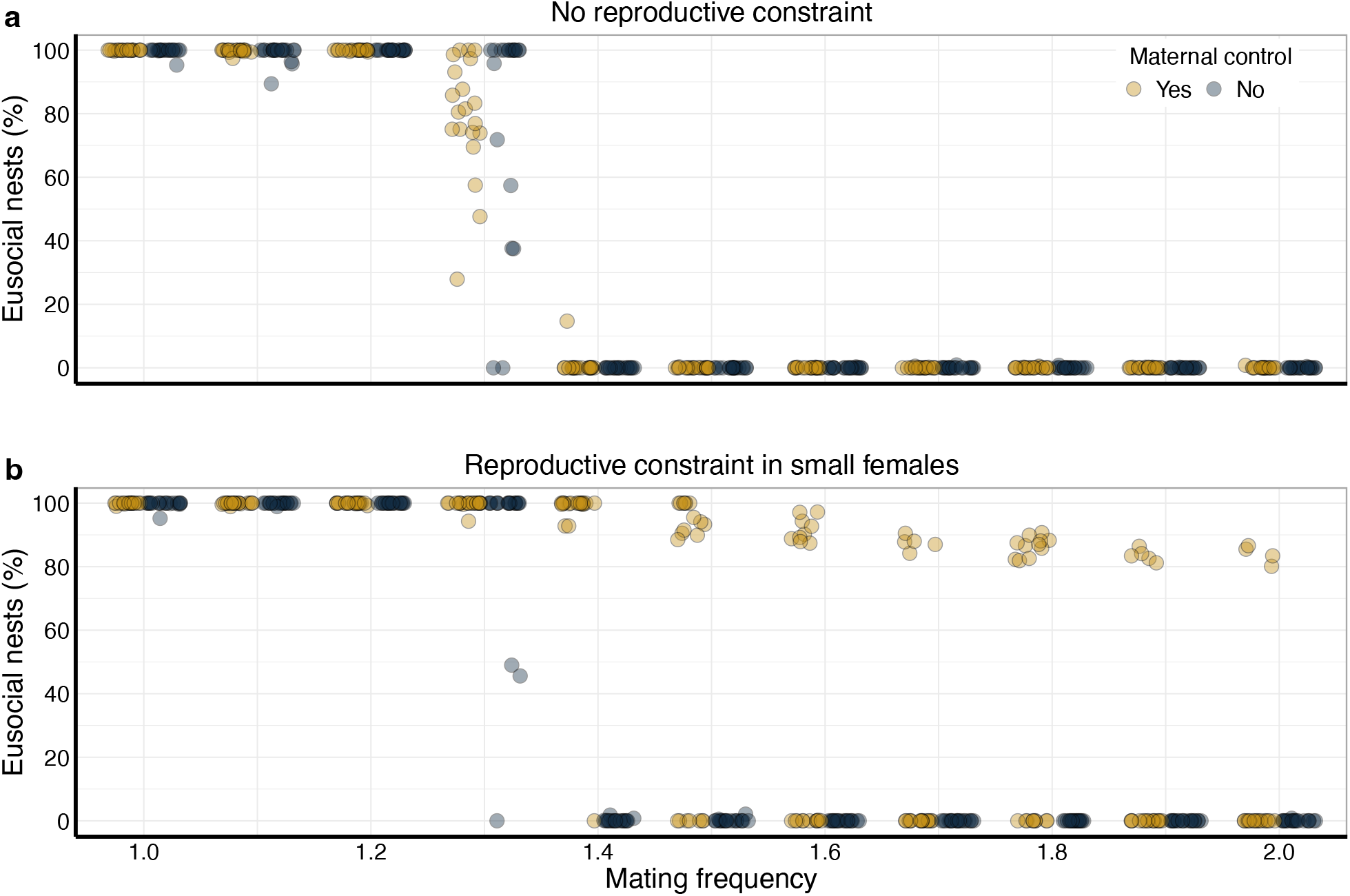
The effect of maternal control over offspring body size and reproductive constraint for small females on the evolution of eusociality. **(a)** Percentage of eusocial nests across different mating frequencies with and without maternal control over offspring body size, assuming no body-size-specific reproductive constraint for small females. **(b)** The same as Fig. 2a, but now assuming a reproductive constraint for small females. Each dot represents the percentage of eusocial nests in the population at the end of a replicate simulation (n = 20 per parameter setting).

### Eusociality evolves despite high levels of polyandry, if small females face a reproductive constraint

Empirical studies on social insects hypothesised that small females might have low breeding success, leading them to forfeit reproduction and become a helper^27–30^. Limited energy reserves in small females might, for instance, hinder their ovarian development, making them less fertile or even incapable of breeding^37–41^, or make them less likely to obtain breeding sites^42,43^. We therefore introduced a body-size-specific reproductive constraint, causing smaller females to gain more resources as helpers than they would as breeders (Fig. 1b). In the absence of maternal control over offspring body size, eusociality only evolves under low levels of polyandry. However, if mothers are capable of controlling offspring body size, then eusociality occurs even under intermediate levels of polyandry (mating frequency of 1.4). At even higher levels of polyandry (mating frequencies of 1.5 and above), social polymorphism emerges with some spring-brood females evolving a disperser- and some a helper-strategy. If the reproductive constraint in small females is stronger, eusociality can evolve at those higher mating frequencies, replacing social polymorphism (Fig. S1-S3). This demonstrates that, with maternal control, eusociality and social polymorphism can evolve under high levels of polyandry.

### Mothers manipulate daughters into helping by imposing a reproductive constraint on them

In order to investigate whether mothers really manipulate their daughters into helping, we ran three different scenarios of the model (all assuming a reproductive constraint for smaller females as in Fig. 2b). First, we prevented the dispersal reaction norm from evolving, causing daughters to become helpers by default (“Monandry + helping by default”). We then obtained an offspring body size distribution that results from the evolved maternal allocation strategy if daughters become helpers (Fig. 3a). Second, we allowed the dispersal probability to evolve under a mating frequency of 1.0 (“Monandry + evolved helping”; Fig. 3b). The body size distributions of helpers obtained under these two scenarios do not differ from one another (mean (0.95 CI): 4.16 (4.11, 4.21) vs. 4.16 (4.10, 4.21); pd = 0.59), demonstrating that maternal manipulation plays no role in the evolution of eusociality under monandry (though see Fig. S4+5). Third, we simulated a case of polyandry (mating frequency 1.8) where eusocial nests only evolved if mothers were able to control offspring body size and small daughters were reproductively constrained (“Polyandry + evolved helping”). Helpers from these simulations were smaller than helpers from the two other scenarios (mean (0.95 CI): 3.00 (2.95, 3.05); pd = 1.00 in comparison with both monandry scenarios), showing that mothers produce smaller offspring in order to manipulate them into helping. Under polyandry, slightly larger females evolve to disperse and breed independently (Fig. 3c), resulting in an S-shaped dispersal reaction norm that underlies the social polymorphism in Fig. 2.

**Fig. 3.**
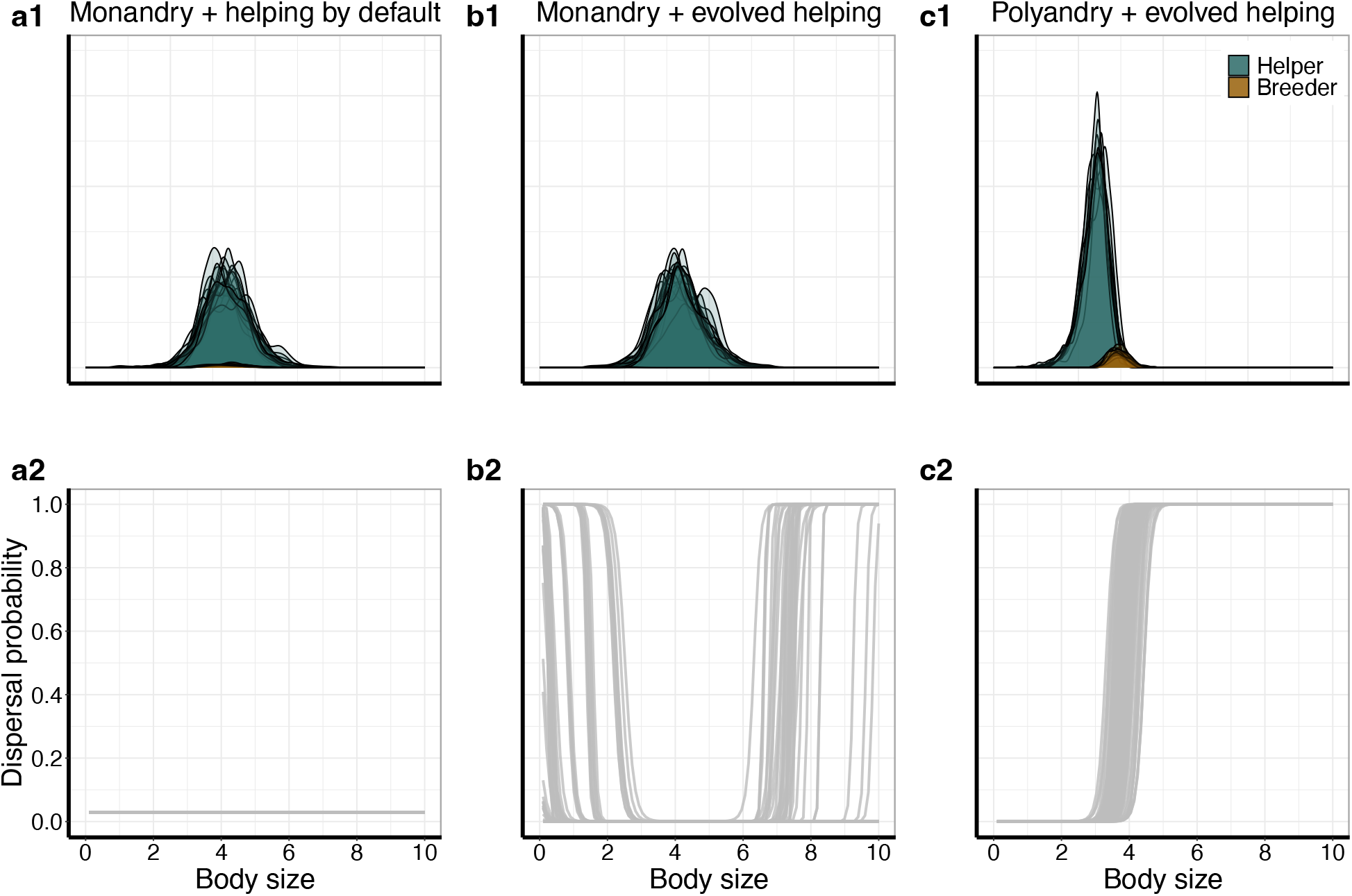
Evolved body size distributions and dispersal reaction norms. **(a1, b1, c1)** Body size distribution of helpers (blue) and breeders (brown). Each distribution represents the body size distribution from one replicate simulation (n = 20 per scenario, but see below). **(a2, b2, c2)** Evolved dispersal reaction norms of 100 random females across replicate simulations, in which eusociality or social polymorphism evolved. **(a1, a2)** Simulations with a mating frequency of 1.0, where dispersal probability was not allowed to evolve and thus females become helpers by default. **(b1, b2)** Simulations with a mating frequency of 1.0 and with evolving dispersal reaction norms. High dispersal probabilities occur at body sizes that are rarely expressed and thus represent cryptic genetic variation. **(c1, c2)** Simulations with a mating frequency of 1.8 and evolving dispersal reaction norms. Only replicates in which social polymorphism evolved are shown (n = 11; n = 9 replicates with solitary breeding not shown here).

## Discussion

We here presented an evolutionary individual-based simulation model to unify kin selection and maternal manipulation explanations for the evolution of eusociality. Maternal control of offspring body size alone does not favour the evolution of eusociality. However, in the realistic scenario^27–30,37–43^, where small females have reduced success at independent breeding, mothers evolve to produce smaller offspring that are manipulated into helping. The offspring, in turn, evolves to help presumably since its inclusive fitness gains from independent breeding are outweighed by those of helping. Similarly, other models also demonstrated that maternal manipulation of offspring behaviour and the offspring’s fitness prospects widens the conditions under which helping and eusociality can evolve^44–47^.

It is currently widely believed that eusociality can only evolve from an ancestor with strict lifetime monogamy. Boomsma^15^, for instance, states that “strict lifetime monogamy […] appears to have been a universally necessary, although not sufficient, condition for allowing the evolution of differentiated eusocial worker castes”. The logic behind this argumentation is intuitive and compelling – strict lifetime monogamy causes relatedness between siblings to be identical to the relatedness between a mother and her offspring. Therefore, the smallest benefit of group living over solitary breeding can tip the balance towards the evolution of eusocial breeding^13–15^. Furthermore, ancestral state reconstruction indicates that the eusocial hymenopterans (ants, bees and wasps) most likely evolved from monogamous solitary ancestors^16^, and in mammals^48^ and birds^49^, cooperative breeding is associated with lower levels of promiscuity than solitary breeding. However, our model demonstrates that while a monogamous mating system is beneficial for the evolution of eusociality, it is not at all a *necessary* condition, since eusociality evolved in our simulations even if the mating frequency was not strictly one. This happens for two reasons. First, male generation overlap in a partially bivoltine life cycle decreases the reproductive value of summer-brood males and thus enables spring-brood females to capitalise on relatedness asymmetries to their sisters vs. brothers from the summer brood due to haplodiploidy (Fig. S6+S7). Quiñones & Pen^24^ showed that this effect is reinforced if mothers can bias sex ratios of the spring and summer brood. In a solitary partially bivoltine life cycle, male generation overlap leads to the evolution of a male-biased spring and a female-biased summer brood. Due to haplodiploidy, females from the spring brood are thus more closely related to their siblings from the summer brood than to their own offspring, leading to the evolution of helping spring-brood females (Fig. S8). Second, in our model, breeding females evolve to impose a fitness cost for independent breeding on their offspring by producing offspring of small body sizes. The offspring consequently evolve to rather help than disperse and breed, even if mothers are multiply mated.

Partial bivoltinism plays a key role in mechanistic explanations for the evolution of eusociality. The diapause ground plan hypothesis suggests that pre-existing morphological or behavioural differences between the two broods of a partially bivoltine life cycle could be co-opted for the evolution of worker- and queen-phenotypes^20,25,26^. Our model combines such more mechanistic explanations with the more classic ultimate explanations for the evolution of eusociality^8^ by demonstrating that phenotypic differences between broods can originate from maternal manipulation of offspring body size. Under high mating frequencies, larger well-nourished females evolve to disperse and breed whereas smaller malnourished females evolve to become helpers. This result matches the prediction of the diapause ground plan hypothesis that a nutrition-dependent developmental switch regulates the production of the worker- and queen-phenotype^20,25,26,50,51^. Accordingly, many eusocial insects with irreversible castes have a nutrition-dependent caste determination system where caste is determined by food obtained during larval development^52–54^.

It is usually assumed that queen-worker dimorphism is an elaboration of eusocial breeding that evolves from a dominance-based breeding system where individuals only temporarily commit to helping^26,55^. Our model demonstrates that phenotypic differences between breeders and helpers can originate from maternal manipulation and thus coincide with the origin of helping behaviours, leading to the evolution of social polymorphism. Such social polymorphism could easily be converted into fully eusocial breeding by a modulation of the partially bivoltine life cycle, which causes breeding females from the spring brood to enter diapause to breed in the next season instead of breeding in the summer. Such an early diapause strategy is indeed observed in some species of partially bivoltine halictine bees, who also exhibit body size differences between helping and breeding females^30,56,57^. The evolution of queen-worker dimorphism might thus coincide with the evolution of helping behaviour and originate from ancestral polyandry and maternal manipulation. Interestingly, in some species of bees (*Halictus rubicundus* and *Ceratina calcarata*), helping females tend to be smaller than breeding females, suggesting a role of maternal manipulation for the evolution of helping behaviours, and both species have been reported to exhibit some degree of polyandry^27,28,57–60^.

Overall, our model presents a realistic scenario for the evolution of eusociality, where eusociality can evolve despite polyandrous mating due to maternal manipulation of offspring body size – a scenario that is realistic because evidence for manipulation, body size differences between breeders and helpers, and polyandrous mating have been found in some social bees. This challenges the current consensus beliefs that monogamous mating is a necessary prerequisite for the evolution of eusociality and that queen-worker dimorphism is a secondary elaboration of eusociality that does not originate at the evolutionary emergence of eusociality.

## Methods

### Model overview

The evolutionary individual-based simulation model follows a population of haplodiploid organisms with a partially bivoltine life cycle over 1,000,000 years. Each simulation starts with *N* identical solitarily breeding females (parameter values in Table 1). Individuals have a body size *X*, which can vary between 0.0 and 10.0 arbitrary units.

**Table 1.** Parameter values used in the simulations, unless stated otherwise.

| Variable | Value | Meaning |
| --- | --- | --- |
| $N$ | 1000 | Population size |
| $f$ | 1.0 | Foundress survival from spring to summer brood |
| $m$ | 0.9 | Male survival from spring to summer brood |
| $s$ | 1.0-2.0 | Mating frequency |
| $a$ | 10, 6.05 | Shaping parameter for resource return function |
| $b$ | 0.5, 1.5 | Minimum body size to gain resources |
| $c$ | 20 | Shaping parameter for resource return function |
| $\mu_X$ | 5.0 | Average default body size |
| $\sigma_X$ | 0.1 | Standard deviation default body size |
| $h$ | 6.2 | Location of logistic larval emergence function |
| $i$ | 7.7 | Steepness of logistic larval emergence function |
| $p$ | 0.05 | Mutation rate |
| $\mu_{\text{mut}}$ | 0.06 | Mutational standard deviation |

### Life cycle

We implemented a partially bivoltine life cycle (Fig. 1a). Each season begins with *N* mated, solitary foundresses, who produce the spring brood. Female offspring from the spring brood disperse to mate with spring-brood males and breed during the summer or they stay at the natal nest to become a helper. Foundresses survive with probability *f* to breed again in the summer. Spring-brood males survive with probability *m* until the summer-brood females emerge. Other parameterisations of *f* and *m* are explored in Fig. S4-S7. The summer brood is produced by surviving foundresses and dispersing female offspring from the spring brood. Female offspring from the summer brood mate with males from the summer brood or surviving males from the spring brood, and subsequently enter hibernation to become the foundresses of the next year. All males die and do not hibernate. During hibernation, population size is re-established by randomly selecting *N* of the hibernating females. During the summer brood, population size can overshoot *N*.

### Dispersal and mating

We assume a flexible natural cubic spline function whose shape is determined by 4 gene values to model the probability of spring-brood females to disperse instead of becoming a helper at the natal nest as a function of their body size (examples in Fig. 1c). Natural cubic splines consist of connected cubic polynomials, which allows them to take highly flexible shapes (though both ends are linear; for details, see Supplement). The simulations start with high dispersal probabilities of 0.97; thus, initially solitary breeding prevails in the population (unless stated otherwise). Females from the spring brood mate during dispersal, and females from the summer brood mate before hibernation. Females store sperm to create their offspring, and do not remate. Across simulations, we varied the mating frequency *s* to manipulate relatedness between siblings from the same nest. A parameter value of 1.0 implies that all females adhere to strict lifetime monandry, whereas values greater than this imply different extents of polyandry. For instance, *s* = 1.5 means that, on average, 50% of females mate once and 50% of females mate twice. The mate(s) is/are selected at random from a global pool of males.

### Resource acquisition

The amount of resources *R* a female can obtain depends on her body size *X*. We selected a function to model this relationship, where very small individuals do not obtain resources and where resource returns level off at large body size. We use the function

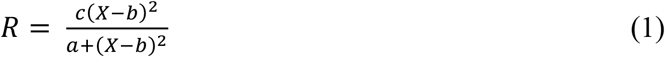

where *a* and *c* are shaping parameters and *b* represents the minimum body size required to gain resources (we set *R* = 0 when *x < b*). In solitary nests, the total resources available are equal to that foraged by the sole female. In eusocial nests, the total resource amount is the sum of the resources obtained by the breeder and those obtained by the helper(s). We focussed on two main parameterisation scenarios of eq. 1: (1) No reproductive constraint for small females – the function is identical for all females, independently of whether they are breeders or helpers (*a* = 10, *b* = 0.5; ‘helper’ function in Fig. 1b); (2) A body-size-specific reproductive constraint for small females – for body sizes of 5.0 and above, all females have the same resource returns for the same body size, but below 5.0, helpers are more efficient in resource acquisition than breeders (function as in (1) for helpers, and breeders with a body size above 5.0, but for breeders smaller than 5.0, *a* = 6.05, *b* = 1.5; ‘breeder’ function in Fig. 1b). Other parameterisations of (2) are explored in Fig. S1-S3.

### Sex allocation and reproduction

Females carry one gene for spring-brood and one gene for summer-brood sex allocation. These genes are associated with numbers that are logistically transformed to determine the proportion of resources invested into males. In the main manuscript, we did not allow resource allocation to the different sexes to evolve and instead fixed it at 50:50 resource allocation to males vs. females. Results with evolving sex allocation are included in Fig. S8. Females produce offspring of both sexes until they run out of resources. We assume that resources invested in offspring translate linearly to offspring body size. We assume some variation in offspring body size which is determined by sampling a normal distribution with mean *μ*_X_ and standard deviation σ_X_. If a female has insufficient resources to create a final offspring, we stochastically decide whether this offspring is still produced by performing a weighted coin flip with a probability given by dividing the remaining resources by the body size of the offspring that is potentially produced. The default body size applies to male offspring from the spring and summer broods, and to female offspring from the summer brood. All offspring have a body-size-dependent emergence probability that determines whether they develop from a larva into an adult. The emergence probability increases with the amount of resources that a larva obtained and is given by

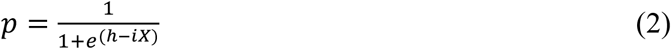

where *f* and *g* are parameters that affect the function’s steepness and location.

### Maternal control of daughter body size

We allow breeding females to have control over resource allocation to female offspring in the spring brood. Females can evolve strategies where their spring broods consist of many small or few large daughters, or where they produce daughters of different sizes. We model this relationship with a flexible natural cubic spline function, whose shape is determined by 4 gene values (details in Supplement). This function determines the body size on the y-axis of the x-th daughter produced by the breeding female. Again, females produce offspring until they run out of resources, and again, it is stochastically decided whether the final daughter, for which insufficient resources exist, is created.

### Genetics and mutation

We assume haplodiploid sex determination; thus, females are diploid and males are haploid. Females carry two sets of 10 genes in total (4 genes determine the shape of the dispersal reaction norm, 4 genes determine the shape of the maternal control reaction norm, and 1 gene each determines spring- and summer-brood sex allocation). Mutations occur by the per-locus mutation rate *p* each meiotic event. If a mutation occurs, the gene value is altered by a value sampled from a normal distribution with mean 0 and standard deviation *μ*_mut_. The genes are assumed to be unlinked and they can recombine freely. Genes are expressed in females; males only function as gene carriers. Females randomly express either the maternal or paternal gene copy on a per-gene basis.

### Model analysis

The model was constructed in C++ and compiled with g++ 11.3.0. We used the Augmented Dickey-Fuller test from the R-package *tseries* v0.10-53^61^ to determine if simulations had reached an evolutionary equilibrium by running the test on the time series of the proportion of eusocial nests in the population. We accepted the assumption of stationarity for p-values less than 0.05. We excluded and reran replicate simulations that were not stationary after 1,000,000 years. We report a nest as being “eusocial” when it had at least one helper in the summer brood, whereas we refer to nests without helpers in the summer brood as being “solitary”. All data analysis and plotting was conducted in R v4.2.1^62^ using the R-packages *tidyverse* v2.0.0^63^, *cowplot* v1.1.1^64^, *stringr* v1.5.0^65^, and *MetBrewer* v.0.2.0^66^. For Fig. 3, we derived probabilities of direction (pd), which represents the certainty that an effect occurs in a particular direction, from Bayesian models implemented with the *brms* v2.20.4^67–69^ package in combination with the MCMC sampler of RStan^70^ and posterior means with the *emmeans* v1.8.8^71^ package (details in Supplement).

## Data availability

All data generated during this study is available under https://doi.org/10.34894/OBFUUV. Preview link for review: https://dataverse.nl/privateurl.xhtml?token=7652fdf9-d10c-470e-a476-5db5ee1e3b00

## Code availability

Simulation code and data analysis scripts are available under https://doi.org/10.34894/OBFUUV. Preview link for review: https://dataverse.nl/privateurl.xhtml?token=7652fdf9-d10c-470e-a476-5db5ee1e3b00

## Acknowledgements

We are grateful to Andrés Quiñones for discussing the model results with us. We thank the Center for Information Technology of the University of Groningen for providing access to the Peregrine high performance computing cluster. JJK was supported by an Adaptive Life grant by the University of Groningen.

## Author Contributions Statement

Conceptualisation: ERB, IP, JJK; Implementation: ERB, IP, JJK; Model analysis: ERB; Writing: ERB, IP, JJK.

