## Supplement for "Maternal manipulation of offspring size can trigger the evolution of eusociality in promiscuous species"

### Strength of the reproductive constraint on small females

In the main manuscript, we present results (1) without a body-size-specific reproductive constraint for small females (all females gain resources according to eq. 1 where $a$ = 10 and $b$ = 0.5) and (2) a small reproductive constraint for small females (all helpers, and breeders larger than 5.0 gain resources as before, but for breeders smaller than 5.0 we set $a$ = 6.05 and $b$ = 1.5). We also investigated the effects of imposing larger reproductive constraints on small females (Fig. S1-S3).

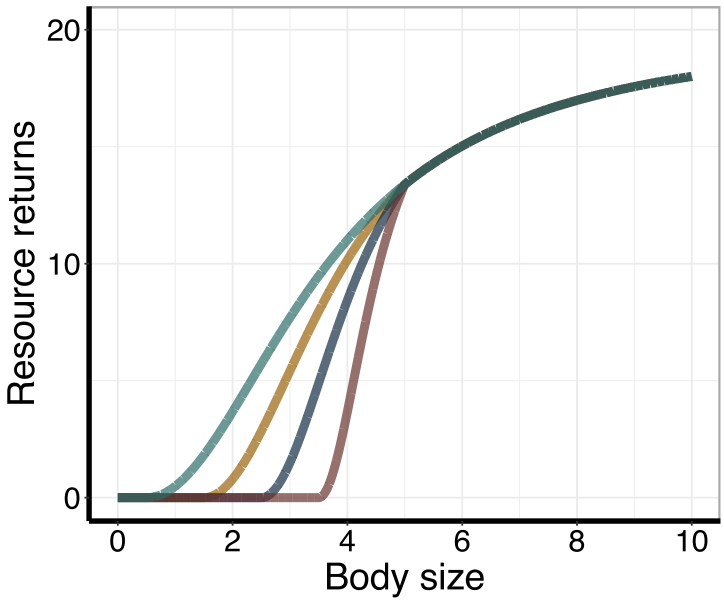


**Fig. S1 | Resource return functions used to investigate the effect of different strengths of reproductive constraint on small females.** Green: helpers in all simulations gain resources according to this function (same as 'helper' function in Fig. 1b; eq. 1 where $a$ = 10 and $b$ = 0.5). Yellow: assuming a small reproductive constraint on small females, breeders gain resources according to this function (same as 'breeder' function in Fig. 1b; eq. 1 where $a$ = 6.05 and $b$ = 1.5 for body sizes less than 5.0). Blue: assuming an intermediate reproductive constraint on small females, breeders gain resources according to this function (eq. 1 where $a$ = 3.09 and $b$ = 2.5 for body sizes less than 5.0). Red: assuming a large reproductive constraint on small females, breeders gain resources according to this function (eq. 1 where $a$ = 1.11 and $b$ = 3.5 for body sizes less than 5.0).

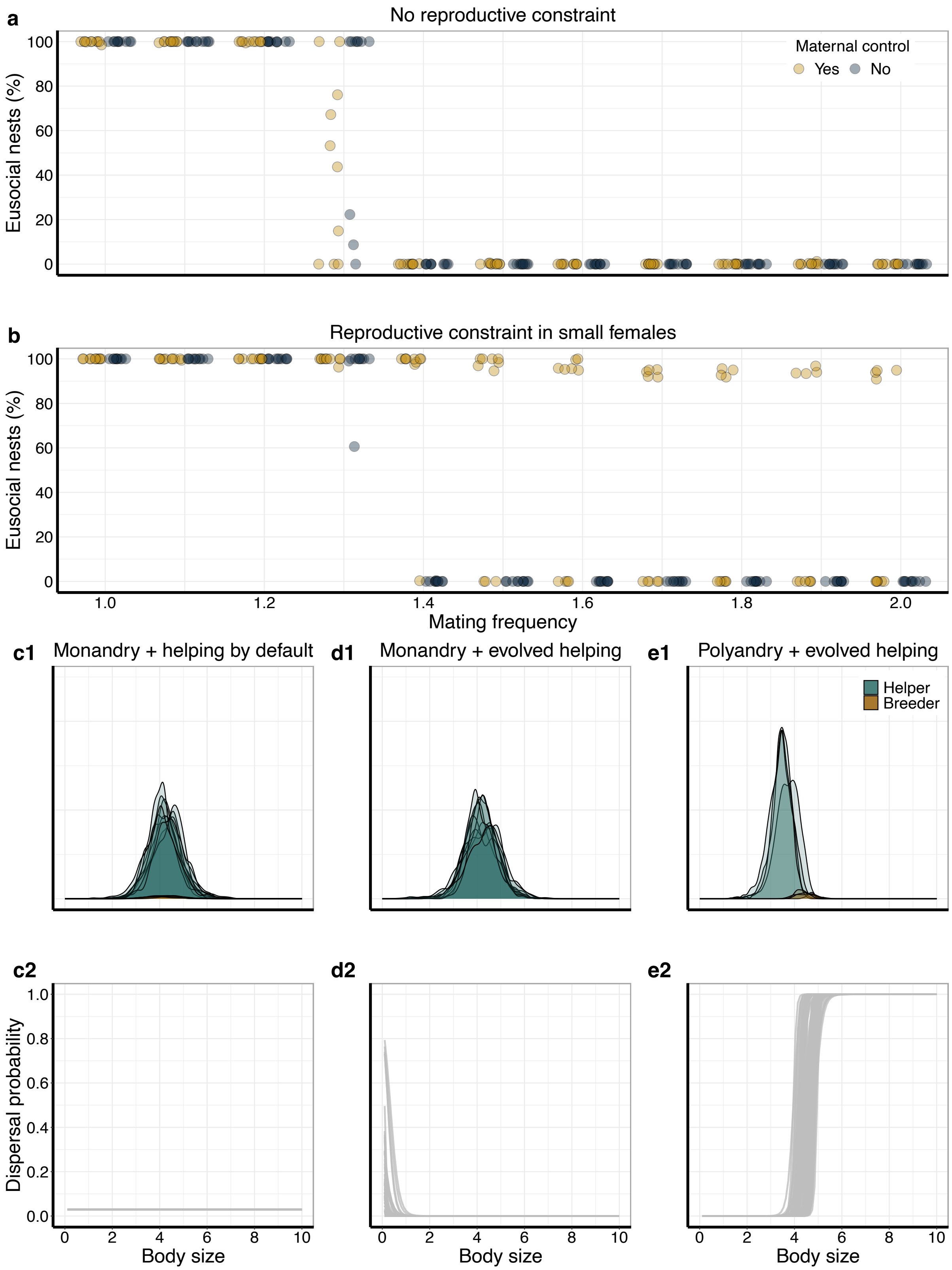

**Fig. S2 | The effect of an intermediate reproductive constraint for small females on the evolution of eusociality (blue function in Fig. S1). (a)** Percentage of eusocial nests across different mating frequencies with and without maternal control over offspring body size, assuming no body-size-specific reproductive constraint for small females. **(b)** The same as Fig. S2a, but now assuming a reproductive constraint for small females. Each dot represents the percentage of eusocial nests in the population at the end of a replicate simulation (n = 10 per parameter setting). **(c1, d1, e1)** Body size distribution of helpers (blue) and breeders (brown) for the three model scenarios described in the main manuscript (Fig. 3). Each distribution represents the body size distribution from one replicate simulation (n = 10 per scenario, but see below). **(c2, d2, e2)** Evolved dispersal reaction norms of 100 random females across replicate simulations, in which eusociality or social polymorphism evolved. **(c1, c2)** Simulations with a mating frequency of 1.0, where dispersal probability was not allowed to evolve and thus females become helpers by default. **(d1, d2)** Simulations with a mating frequency of 1.0 and with evolving dispersal reaction norms. High dispersal probabilities occur at body sizes that are rarely expressed and thus represent cryptic genetic variation. **(e1, e2)** Simulations with a mating frequency of 1.8 and evolving dispersal reaction norms. Only replicates in which social polymorphism evolved are shown (n = 4; n = 6 replicates with solitary breeding not shown here).

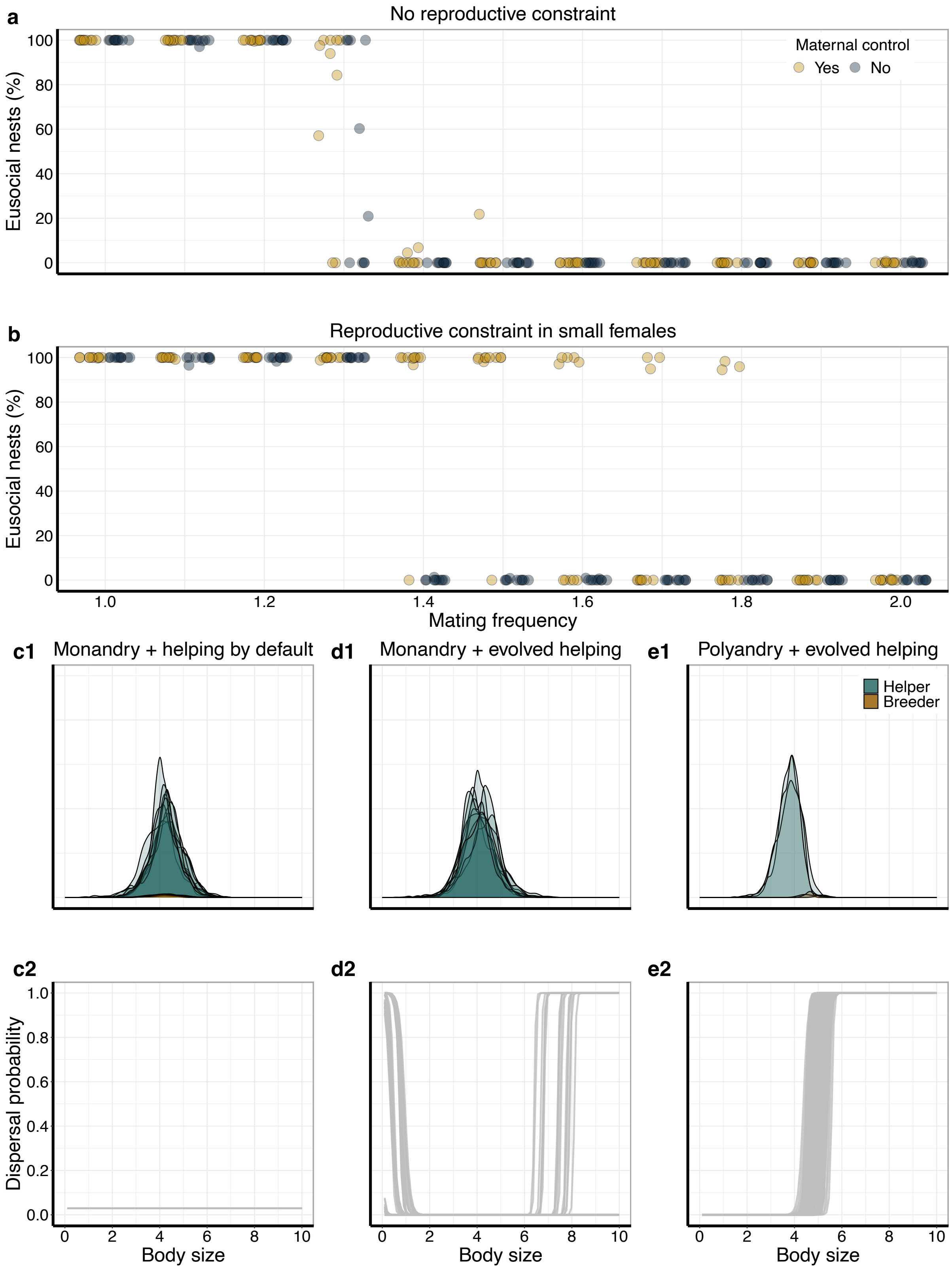


**Fig. S3 | The effect of a large reproductive constraint for small females on the evolution of eusociality (red function in Fig. S1).** Figure conventions as in Fig. S2. **(e1, e2)** n = 3; n = 7 replicates with solitary breeding not shown here.

**The effect of foundress survival on the evolution of eusociality**

In the results in the main manuscript, foundresses have a high probability $f = 1.0$ to survive from the spring to the summer brood. We also investigated the effects of intermediate ($f = 0.85$; Fig. S4) and low ($f = 0.5$; Fig. S5) foundress survival. In the model, we assume that helping females gain no fitness through reproduction or helping when the foundress in the nest, where they help, dies. These conditions make the evolution of philopatry and helping the least favourable. Thus, our assumptions were the most conservative for the evolution of eusociality. Low foundress survival therefore imposes a cost upon philopatric, helping females because, if the foundress dies, they can neither reproduce nor help raise offspring. Under intermediate rates of foundress survival, helping does not evolve in the absence of a reproductive constraint for helpers (Fig. S4a). However, if mothers can control offspring body size and if small females face a reproductive constraint, then mothers can manipulate their daughters into helping (Fig. S4b-e2). In this case, helping evolves by maternal manipulation, even under monandrous mating. If foundress survival, however, is too low, then helping does not evolve since it is more beneficial for mothers to produce dispersing offspring that breed rather than helpers (Fig. S5).

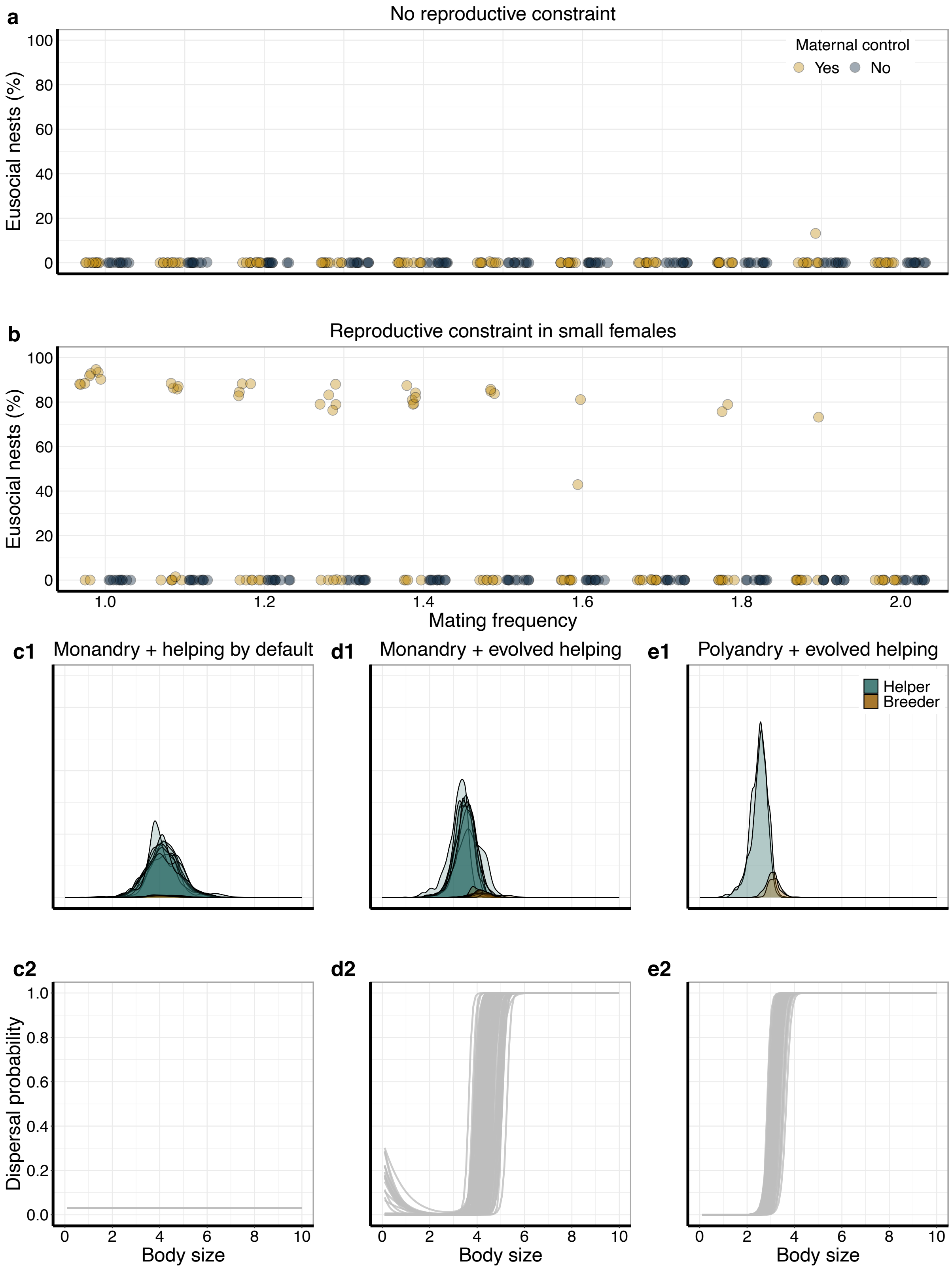


**Fig. S4 | The effect of intermediate foundress survival (*f* = 0.85) on the evolution of eusociality.** Figure conventions as in Fig. S2. **(e1, e2)** n = 2; n = 8 replicates with solitary breeding not shown here.

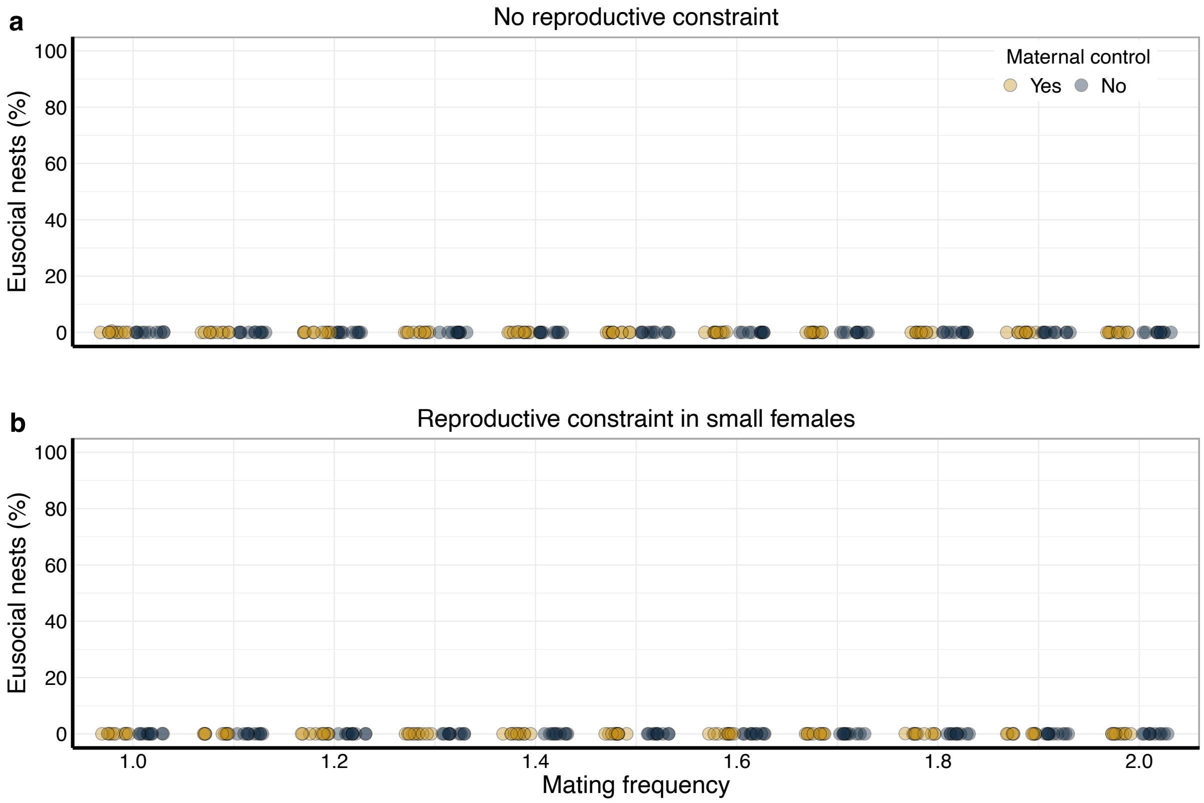


**Fig. S5 | The effect of low foundress survival (*f* = 0.5) on the evolution of eusociality.** Figure conventions as in Fig. S2a+b.

### The effect of male survival on the evolution of eusociality

In the results in the main manuscript, all male offspring have a high probability $m$ = 0.9 to survive from the spring to the summer brood. We also investigated the effects of intermediate ($m$ = 0.5; Fig. S6) and low ($m$ = 0.1; Fig. S7) male survival probabilities. We recover results consistent with those of Quiñones & Pen[1] showing that higher male survival widens the conditions under which eusociality can evolve. This is because high male survival decreases the reproductive value of summer-brood males which enables spring-brood females to capitalise on relatedness asymmetries to their sisters vs. brothers from the summer brood due to haplodiploidy.

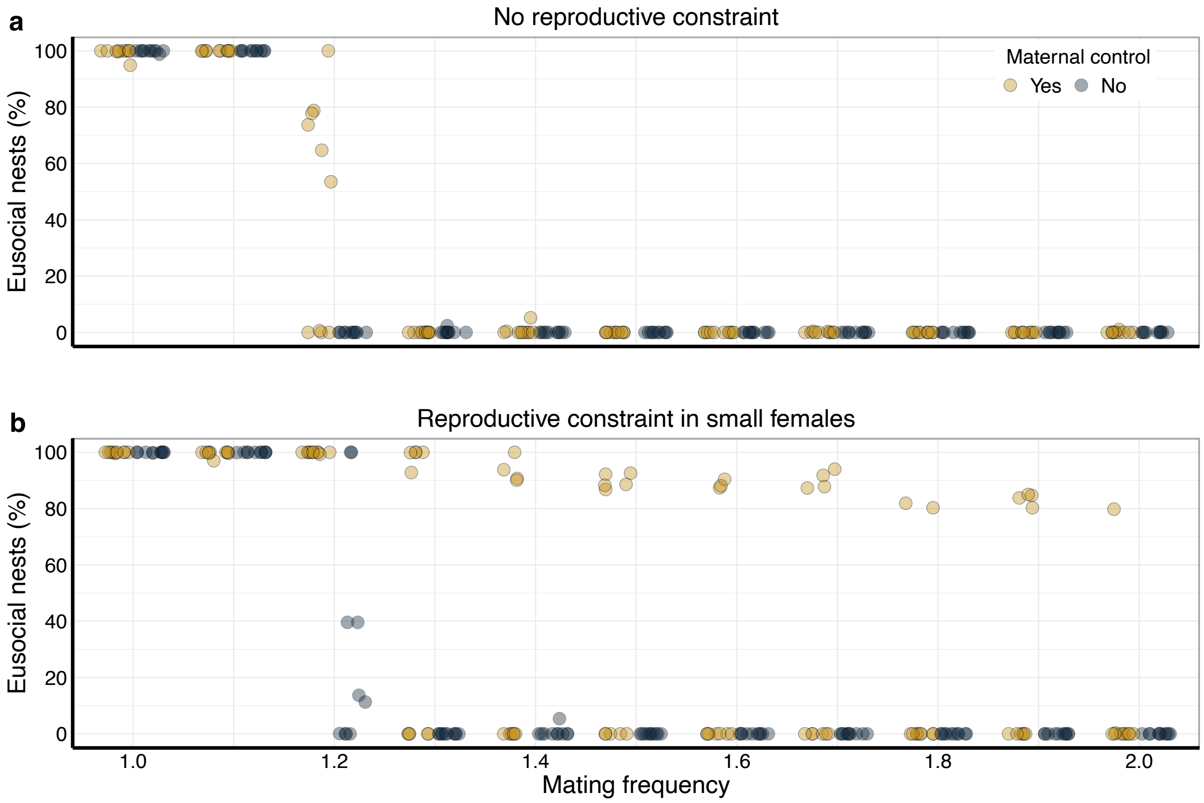


**Fig. S6 | The effect of intermediate male survival probability (*m* = 0.5) on the evolution of eusociality.** Figure conventions as in Fig. S2a+b.

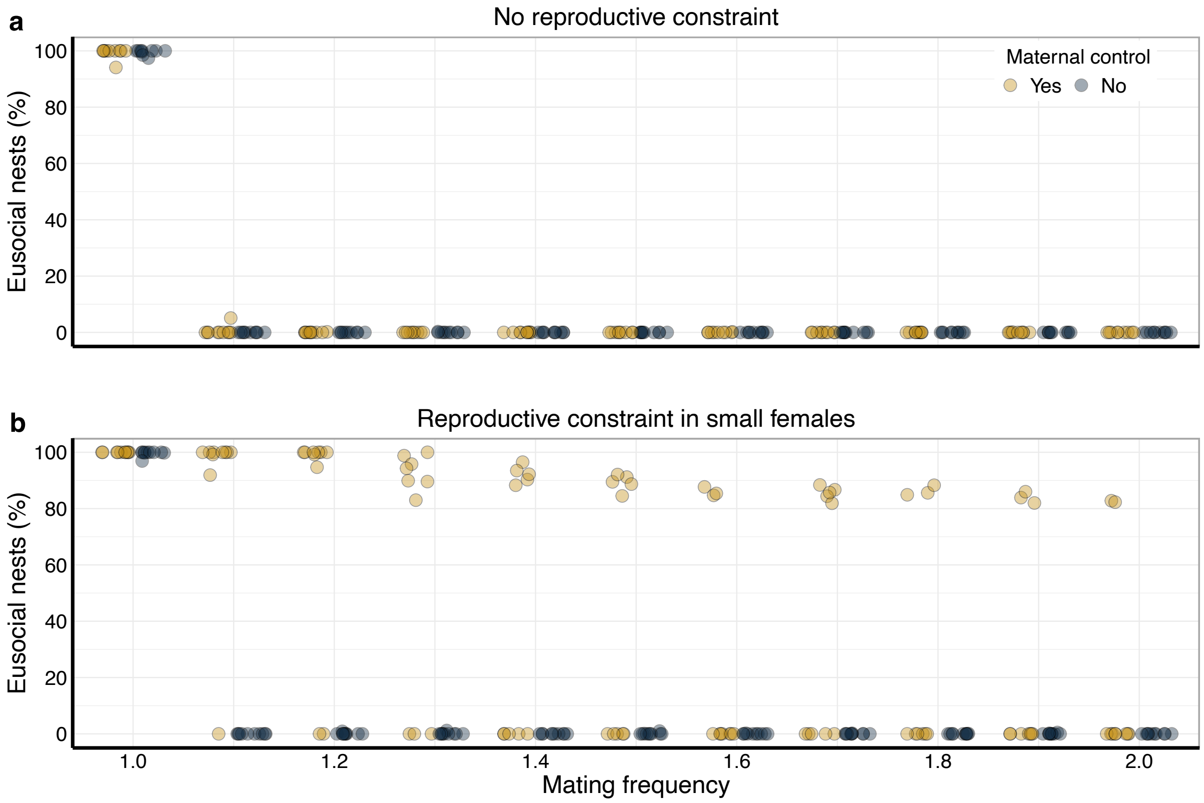


**Fig. S7 | The effect of low male survival probability (*m* = 0.1) on the evolution of eusociality.** Figure conventions as in Fig. S2a+b.

**The effect of evolving sex allocation on the evolution of eusociality**

In the main manuscript, sex allocation in the spring and summer brood are both fixed to 50% allocation to daughters and 50% allocation to sons. We here present additional results where sex allocation was allowed to evolve. As in Quiñones & Pen[1], we first let sex allocation evolve in a population that is solitary, i.e., individuals disperse by default and cannot evolve helping (first 200,000 years of 1,000,000 years simulation). After this 'solitary phase', in which sex allocation evolves to be male biased in the first and female biased in the second brood, we enable the evolution of the dispersal probability and thus of helping. This leads to the evolution of a female-biased first brood and approx. even sex allocation in the second brood. Our results are in line with the results of Quiñones & Pen[1] in that evolving sex allocation widens the conditions under which eusocial nests occur (Fig. S8).

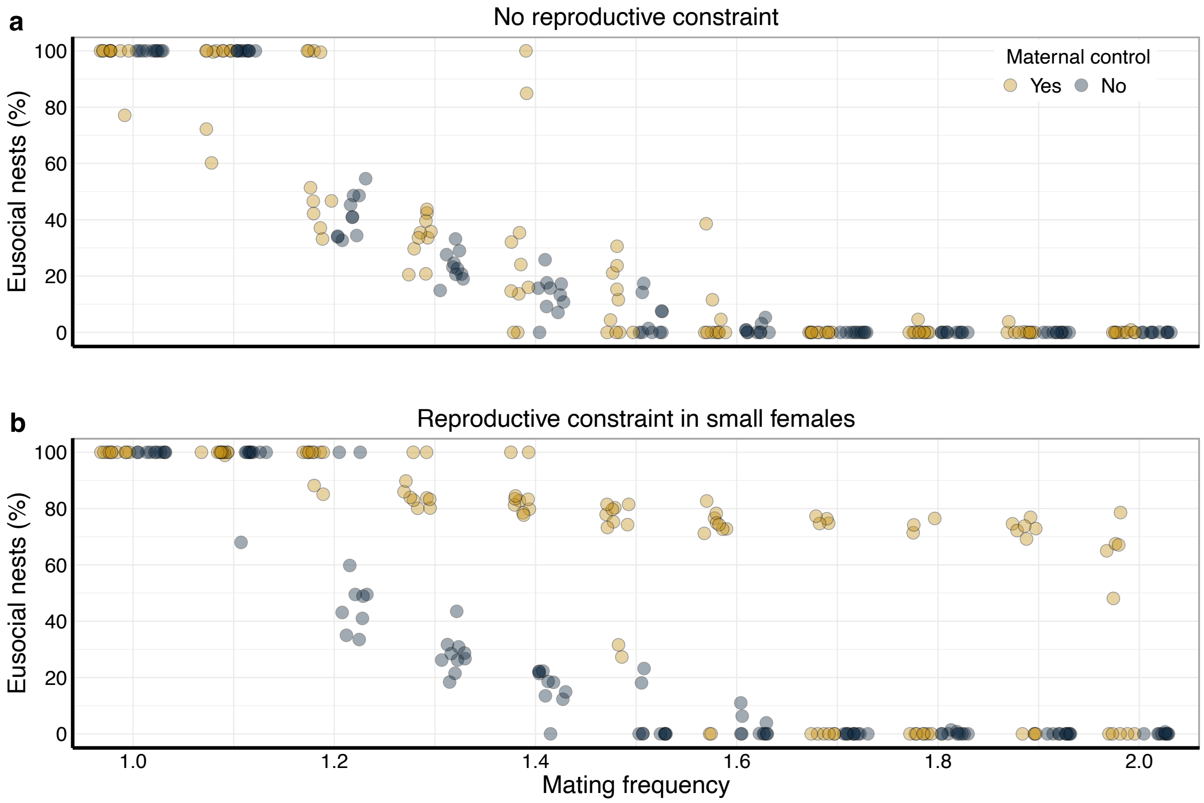


**Fig. S8 | The effect of evolving sex allocation on the evolution of eusociality.** Figure conventions as in Fig. S2a+b.

**Modelling reaction norms with natural cubic spline functions**

We model flexible smooth reaction norms with natural cubic spline functions as done by Lagos-Oviedo et al.[2]. These functions consist of connected cubic polynomials but they are linear on both tail ends. We follow the definition of a natural (or restricted) cubic spline function by Harrell[3]. A natural cubic spline function has $k$ knots, which are the connection locations of the cubic polynomials with respect to $x$ (in the main manuscript, we set $k = 4$). The knot locations of the $k$ knots are $t_1, \ldots, t_k$. The natural cubic spline function, consisting of the basis functions $B_i$, is then given by

$$f(x) = \beta_0 B_0 + \beta_1 B_1 + \beta_2 B_2 + \cdots + \beta_{k-1} B_{k-1}, \qquad (1)$$

where $B_0 = 1$ and $B_1 = x$. The remaining terms $B_2, \ldots, B_{k-1}$ are calculated by iterating over $j = 1, \ldots, k - 2$ according to

$$B_{j+1} = (x - t_j)_+^3 - \frac{(x-t_{k-1})_+^3 (t_k - t_j)}{t_k - t_{k-1}} + \frac{(x-t_k)_+^3 (t_{k-1} - t_j)}{t_k - t_{k-1}}, \qquad (2)$$

where $(\ldots)_+$ indicates that a term is set to 0 when it evaluates to a number below 0. The natural cubic spline function can also be written in matrix notation.

$$f(x_i) = \begin{bmatrix} B_0(x_1) & B_1(x_1) & B_2(x_1) & \cdots & B_{k-1}(x_1) \\ B_0(x_2) & B_1(x_2) & B_2(x_2) & \cdots & B_{k-1}(x_2) \\ \vdots & \vdots & \vdots & \ddots & \vdots \\ B_0(x_n) & B_1(x_n) & B_2(x_n) & \cdots & B_{k-1}(x_n) \end{bmatrix}_i \begin{bmatrix} \beta_0 \\ \beta_1 \\ \beta_2 \\ \vdots \\ \beta_{k-1} \end{bmatrix} \quad (3)$$

Here, the matrix is a so-called basis matrix of the natural cubic spline function. The column vector $\beta$ contains the parameters $\beta_0, \dots, \beta_{k-1}$ for the natural cubic spline function. The basis matrix subscript $i$ signifies that the $i$-th row of the basis matrix is used for the multiplication with the column vector $\beta$. In our simulations, we assume that the knot locations are evenly distributed within the range of $x$. The basis matrix is precalculated on equally-spaced intervals at locations $x_i$ at initialisation for a given resolution of the natural cubic spline function for a given range of $x$ (in the main manuscript, we set the resolution to 100). In the case of the reaction norm for dispersal, we set the range of $x$ between $X_{\min} > 0.0$ and $X_{\max} = 10.0$. The natural cubic spline function parameters in column vector $\beta$ are the evolving gene values. In the reaction norm for dispersal, we initialised $\beta_0 = 3.5$ and all further $\beta$ with 0. Consequently, all females have an initial dispersal probability of 0.97 (from a logistic transformation of 3.5), regardless of their body size $X$. When evaluating the reaction norm for individual female, we identify the two closest prespecified $x_i$ to the body size $X$ of the individual and select one of these. Subsequently, we multiply the column vector $\beta$ with the $i$-th row of the basis matrix. This yields a single value, which is the phenotypic value (in this case, the dispersal propensity) for a particular value of $x$. To arrive at a dispersal probability, we logistically transform the dispersal propensity.

The same method is also used for the function that determines maternal resource allocation to offspring. However, in this case we set the range of $x$ between $D_{min} = 1$ and $D_{max} = 9$. Evaluating the function at $D_j$ yields the body size of the $j$-th daughter. The column vector $\beta$ again contains the evolving gene values. We initialised $\beta_0 = 5.0$ and all further $\beta$ with 0. Consequently, all daughters have an initial body size of 5.0. We transform the result $v$ from the evaluation of the spline function to keep it within the range of body sizes from 0.0 to 10.0 by

$$X = \frac{10}{1+e^{(-v)}} \quad (4)$$

**Details on the Bayesian statistical analysis**

We statistically compared the body sizes of individuals from the three model scenarios in Fig. 3 of the main manuscript using Bayesian models with the *brms*[4–6] package in combination with the MCMC sampler of RStan[7]. We predicted body size (response variable) by the interaction of model scenario and the type of the individual (breeder vs. helper) while also incorporating a random effect of the replicate simulation from which the individual originated. We allowed for variances to differ between groups. We set the response type to 'gaussian'. We used weakly informative Gaussian priors[8,9] with mean = 0 and SD = 10 for intercepts and SD = 1 for the other regression coefficients for the fixed effect parameters. The random effect parameters used the default priors of *brms* (t-density with df = 3 for standard deviations). We ran the model for a subset of 10,000 randomly sampled individuals to decrease computation time. We ran the model with four chains and discarded the first 1000 warm-up iterations, followed by 3000 sampling iterations, resulting in 12000 posterior samples. We monitored proper mixing of chains with trace plots and convergence of chains by verifying that all $\hat{R} = 1.00$. We used the *pp_check*-function from the *brms* package to evaluate model fit by inspecting posterior predictive checks. We obtained predictions for group-specific means and 0.95CI using the *emmeans*[10] package. We calculated the probability of direction (pd), which represents the certainty that an effect occurs in a particular direction, from the overlap of group-specific posterior distributions.
